# An agent based model of intracellular ice formation and propagation in small tissues

**DOI:** 10.1101/2022.03.22.485258

**Authors:** Fatemeh Amiri, James D. Benson

**Affiliations:** Department of Biology, University of Saskatchewan, Saskatoon, SK, Canada

## Abstract

Successful cryopreservation of tissues and organs would be a critical tool to accelerate drug discovery and facilitate myriad life saving and quality of life improving medical interventions. Unfortunately success in tissue cryopreservation is quite limited, and there have been no reports of successful long term organ cryopreservation. One principal challenge of tissue and organ cryopreservation is the propagation of damaging intracellular ice. Understanding the probability that cells in tissues form ice under a given cryopreservation protocol would greatly accelerate protocol design, enabling rational model-based decisions of all aspects of the cryopreservation procedure. Established models of intracellular ice formation (IIF) in individual cells have previously been extended to small linear (one-cell-wide) arrays to establish the theory of intercellular ice propagation in tissues. However these small-scale lattice-based tissue ice propagation models have not been extended to more realistic tissue structures, and do not account for intercellular forces that arise from the expansion water into ice that may cause mechanical disruption of tissue structures during freezing. To address these shortcomings, here we present the development and validation of a lattice-free agent-based stochastic model of ice formation and propagation in small tissues. We validate our Monte Carlo model against Markov chain models in the linear two-cell and four-cell arrays presented in the literature, as well as against new Markov chain results for 2 × 2 arrays. Moreover we expand the existing model to account for the solidification of water into ice in cells. We then use literature data to inform a model of ice propagation in hepatocyte disks, spheroids, and tissue slabs. Our model aligns well with previously reported experiments, and demonstrates that the mechanical effects of individual cells freezing can be captured.

**Author summary:** The widespread ability to successfully store, or cryopreserve, tissues and organs in liquid nitrogen temperatures would be game changing for human and animal medicine and drug discovery. However, success is limited to a select number of small tissues, and no organs can currently be stored in a frozen or solid state and survive thawing. One major contributor to damage during this process is the formation of intracellular ice, and its associated cell level damage. This ice formation is complicated in tissues by the number of intercellular connections facilitating intercellular ice propagation. Previous researchers have developed and experimentally validated simple one dimensional models of ice propagation in tissues, but these fail to capture complex tissue geometries, and have many fewer intercellular connections compared to three dimensional tissues. In this paper, we adopt previous models of ice formation and propagation to a model capable of capturing arbitrary cell orientations in three dimensions, allowing for realistic tissue structures to be modelled. We validated this tool on simple models and with experimental data, and then test it on three structures made of digital liver cells: disks, spheroids, and slabs. We show that we can capture new information about the interaction of cooling the tissue, the formation of intracellular ice, the movement of ice from one cell to another, and the mechanical disruption that occurs during this process. This allows for novel insights into a mechanism of damage during cryopreservation that is cooling rate and tissue structure dependent.

## Introduction

The need for cryopreservation grows alongside the rapidly advancing science and medicine it facilitates. The application of cryobiology in science and medicine has been a great success and includes preserving blood, gametes, and cell therapeutics, but also cartilage, heart valves, corneas, and other small tissues [1–8]. Unfortunately, most tissues and all organs cannot currently be successfully cryopreserved [9]. This is not due to lack of either medical or financial demand. For example, ballooning costs in drug discovery are exacerbated by poor availability of human tissue models that provide the most relevant clinical data, and cryobanked human tissue slices could provide within patient comparisons of the specific actions of propsective drug compounds on tissues of interest. On a larger spatial scale, the demand for heart transplantation in the United States has been estimated to be more than ten times the heart transplant waiting list [9], and organ cryopreservation would make the limiting challenges of donor-recipient matching, organ transport, and even surgery scheduling much simpler. Unfortunately, there are only a few preliminary and limited reports of successful organ preservation in the absence of ice for the period of less than a week [10]. In short, there is a worldwide need for improvement in the recovery of tissues and organs after cryopreservation.

Physical injury of biological cells during freezing can be caused by ice crystalization, inter- and intra-cellular forces that arise from the expansion water into ice, or the extreme dehydration and associated high salt concentrations due to the formation of extracellular ice that provides a driving force for both dehydration of the cell and for the nucleation and growth of damaging intracellular ice [11]. A number of theoretical methods have been developed to improve the design of cryopreservation protocols and minimize the damage created by physical injury. Among very early models applied towards cryopreservation is the water transport model that was developed by Mazur [12, 13] and later on modified by Levin [14]. This model captures the dynamics of the water volume of the cells during cooling at rates where the extracellular medium is not supercooled. Mazur proposed that this model was useful because damaging intracellular ice formation (IIF) could be avoided by cooling cells slowly enough so that the osmotic dehydration allows the intracellular milieux to remain in chemical potential equilibrium with the external environment. In fact, the definition of “slowly enough” could be tested theoretically using the model, and this was borne out in experiments that would go on to support one side of Mazur’s “two factor hypothesis” [15]. Mazur concluded that the alignment with experiments suggested that cells with less than 2 °C of theoretically predicted supercooling were not likely to have intracellular ice formation [15].

This work to predict intracellular ice formation was refined significantly with a statistical approach for predicting the likelihood of intracellular ice, introduced by Pitt [16], that had some success in approximating the probability of intracellular ice formation in different cell types. However, Pitt’s model does not provide information about the mechanisms of ice nucleation in the supercooled cytoplasm. Following Pitt, Toner *et al*. developed a physicochemical theory of heterogeneous nucleation in cells based on a thermodynamic model of ice nucleation, coupled with Mazur’s model of cell dehydration during freezing [17]. Finally, Karlsson *et al*. [18, 19] further refined Toner’s model by adding diffusion limited growth that depended on intracellular viscosity, an important consideration in the multimolar cryoprotectant concentrations that exist in subzero temperatures during cryopreservation.

Although these models were successfully implemented in individual cells [20, 21], cryomicroscopic observations indicated that the probability of IIF in cells in cell monolayers, colonies, and tissues is higher than the probability of IIF in suspended cells of the same type [22]. These findings suggest that cell-cell interaction plays a major role in intercellular ice nucleation. To address this with a mathematical model, Irimia and Karlson extended the isolated cell IIF models to account for the propagation of IIF from one cell to a neighbouring cell using a Markov process for a simple two cell construct, and successfullly validated their theory with experiment [23]. Ostensibly, this was the first cell-based model of ice propagation in tissues, and Irimia and Karlsson then extended this two-cell model and experiments to linear four-cell arrays [24]. In these cell-based models, the Markov chain and the resultant ordinary differential equation (ODE) were solved for the probabilities of each IIF outcome of the tissue. Because the state space of the generator matrix in Markov chain model grows exponentially with the number of cells [25], a numerical Monte Carlo approach was also implemented and tested in linear arrays of up to 100 cells. Finally, work extending to a two dimensional array was presented in an otherwise unpublished Master’s thesis [26], and an exploration of ice propagation in a somewhat unrealistic and spherically symmetric tissue [27, 28].

These small-scale lattice-based tissue ice propagation models have not been extended to more realistic tissue structures, and do not account for intercellular forces that arise from the expansion water into ice that may cause mechanical disruption during freezing. Moreover, the number of intercellular connections are three to four times higher in a three dimensional tissue, and as such the intercellular ice propagation rates and the importance of the types of connections governed by cell type and structures will play an important and unexplored role in the ice dynamics in these tissues. Additionally, as some cell types and structures may tolerate intracellular ice, while others do not, the location of each individual cell type and the likelihood that intracellular ice will cause damage should allow modeling of whether tissues will lose viability during any specific cooling protocol. Finally, the complex interaction of heat and mass transfer and the resultant intracellular ice formation in large tissues plays a critical role during freezing, and is coupled with cell and tissue level osmotic response or temperature change, that, for example, is delayed at the centre of a tissue compared to that at the tissue boundary [14, 29].

While Karlsson, Irimia, and Sumpter stopped at one and two dimensional structures for their cell-based models, there have been a number of other attempts to capture the three dimensional propagation of ice in tissues and organs. These have used continuum models such as the Johnsson-Avrami-Mehl-Kolmagorov model that generalizes the classic Stefan problem of the propagation of a phase transition through a homogenous material [30]. In the case of cryobiology, these models have been mostly applied to understanding the extents of ice balls formed during ablative cryosurgery [31, 32]. They were also applied and compared with the cell-based models by Sumpter [26]. While thermodynamically correct, these continuum models are challenging to implement in homogenous tissues, and become even moreso in the spatially heterogenous tissues relevant to biology. They also lack the cell-level detail to elucidate the impacts of ice propagation rates and damage on key cell types and tissue substructures. Therefore, while continuum models of ice propagation give a good overview of the rates at which ice may form throughout a tissue, they lack the precision and detail needed to understand why some parts of tissues survive and others do not under any particular freezing protocol.

Agent based models have been used in computer science and social science to study self-organizing computer programs, robots, and individuals [33–36]. They have been applied in biological systems to model capsule shaped bacteria [37], molecular systems in biology [38], and cancer and tumor growth and mechanics [39, 40]. Agent based models are generally characterized by self-determining objects called agents equipped with protocols for inter-agent interaction, such as the ability to recognize and distinguish the features of other agents [41].

In this manuscript we couple the cell-based multicellular stochastic model introduced in [24] with a modern agent-based method to model lattice-free ice propagation in tissues and account for the solidification of water into ice in cells. In our approach, each cell in the tissue is considered as an agent using the open source software PhysiCell [40], a multicellular system simulator which is designed to model tissues involving many interacting cells in multi-substrate 3D-microenvironments. This platform is C++ code-based and has only two external dependencies, pugixml [42] and BioFVM [43] and has been parallelized in OpenMP to make use of multi-core computers. The software has been applied to cancer modelling and treatment, including anti-cancer biorobots and cancer heterogeneity and immune response, and has the ability to simulate cell movement, growth, division, interaction, and death. We also take advantage of the continuum model by coupling the thermodynamic theory of ice formation in individual cells introduced in [17] and extended it to tissue size by using the stochastic model introduced in [24] to account for ice propagation in the neighbour cells. Finally, we take advantage of the mechanical self-organizing property of this model and implement the effects of ice formation and expansion in a relatively large tissue in a lattice-free setting. We show here that agent based models are ideal to capture the intercellular propagation of ice in tissues, and allow for the formulation of biologically realistic structures that may influence the rate of propagation.

The paper is organized as follows. In the Mathematical Models section, we introduce the stochastic model presented in [24] and its implementation and application with agent-based modeling. In this section, we study a example of a four cell structure. In the Water Transport Model section, the water transport model is presented in order to model the mechanism of ice formation inside biological cells during freezing based on physicochemical model introduced in [17]. In this section, we also present a model for nondimensional time as a function of real time and temperature. To implement our model in simple tissues, we choose to study hepatocyte disks and spheroids, as this tissue is comprised of a single cell type that has existing independent IIF data and parameters already experimentally determined in individual cells. Thus, in the Results and Discussion section, we first fit the nondimensional time to previously published experimental results for individual rat hepatocyte cells. We then use these new parameters to inform the ice propagation agent based model in this small three dimensional tissue. Next we use the hepatocyte data to explore ice propagation through a larger tissue, and briefly study the impact of water-to-ice volume expansion on tissue structure. Some concluding remarks are given in at the end.

## Mathematical Models

### Stochastic Model of Intracellular Ice Formation

As water turns into ice the large number of random possibilities associated with hydrogen bond rearrangements between individual water molecules means that ice formation is a stochastic event [44]. To account for this in the case of individual cells undergoing solidification, Karlsson [18, 19], following the work of Toner [17] and Pitt [16], introduced a diffusion-limited stochastic model describing the probability of intracellular ice formation given key submodels that include intracellular viscosity and osmolality as a function of temperature. This model was applied to a number of cell types, including oocytes [18, 45] and hepatocytes [19]. To extend this to include the possibility that ice preferentially propagates from a cell with ice to a cell without ice, Irimia and Karlsson [23, 24] developed a multicellular stochastic model, where the mechanisms of IIF were described by two independent stochastic processes: *J*^i^, the class of all mechanisms that cause IIF independent of their neighbouring cells’ states, and *J*^p^, the total rate of intracellular ice propagation across the interface between a frozen and unfrozen cells. Hence, the total rate of IIF in a unfrozen cell *j* is the sum of the rates associated with these two independent stochastic functions. Assume that cell *j* has *k_j_* frozen cell neighbours, then assuming intracellular propagation rates are not cell or position dependent we have

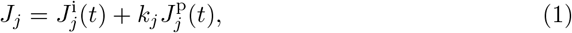

where *J_j_* can be considered to be all possible observations of IIF per unit time per cell. The domain of the *j*th cell’s neighbourhood is set to be the cells which have a centre within a ball of radius *R* around the centre of the *j*th cell, as indicated in Fig 1. Then, the probability that an unfrozen cell *j* freezes in a time interval Δ*t* can be described by the Poisson process

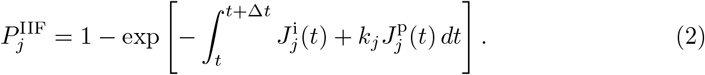

To non-dimensionalize the above equation and simplify the computational model, a nondimensional time *τ* can be introduced [23],

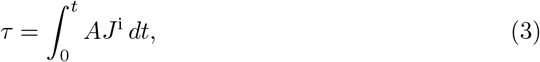

where *A* is the surface area of the cell. Also, a non-dimensional ice propagation rate can be defined as

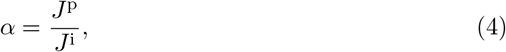

where *α* can be considered constant if the cooling rate fast enough so that cell dehydration is insignificant. Note that *α* controls the propagation speed: the larger *α*, the faster the ice propagates from neighbour to neighbour.

**Fig 1.**
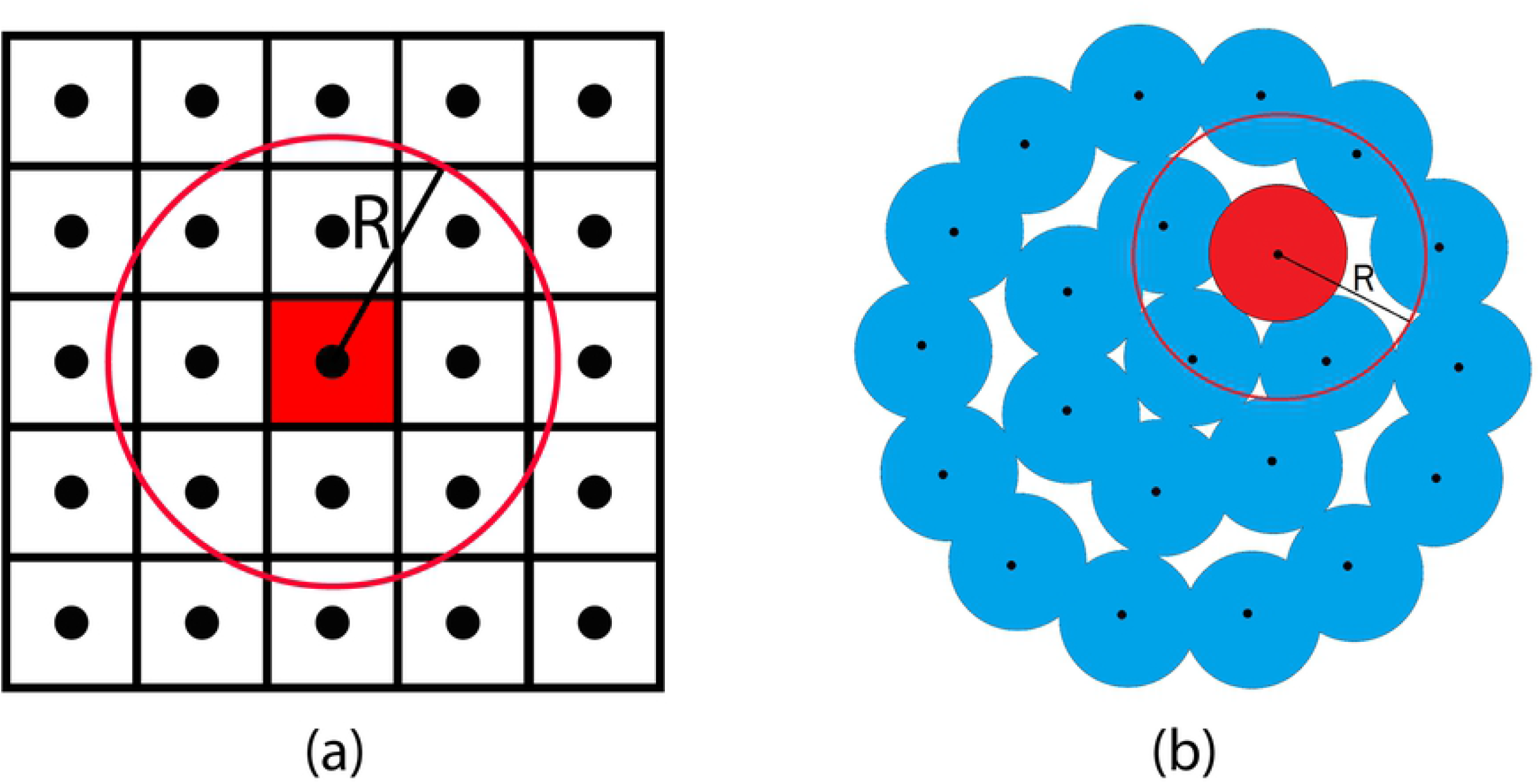
Geometry of neighbourhoods. Cells which have a centre within a ball of radius *R* and the same centre of the cell of interest *j*, here is indicated in red, consider to be the neighbour cells. (a) A “square” cell configuration where the number of neighbours for a central cell *j*, *k_j_* = 8. (b) A two dimensional “single layer” configuration of round cells, the number of neighbours for the red cell *j* is *k_j_* = 4.

These nondimensional parameters make the model independent of the physical mechanisms underlying the interaction-independent IIF process, or the functional form of *J*^i^(*t*). The nondimensional parameter *α* is discussed in more details in [23, 24]. In reality, tissues are heterogeneous and may thus have different *J*^i^ and *α*. In fact, there are only a handful of cell types with measured *J*^i^ and *α* parameters. This is an active area of research in our group.

Finally, as the number of cell neighbours determines the relative paths for ice to propagate from one cell to another, it is important to be clear on the local neighbours of individual cells. Here we define the neighbourhood of a cell *j* to be other cells that have a centre within a ball of radius *R* away from cell *j*, as indicated in Fig 1. Note that the number of neighbour cells depends on the size of radius *R*, and the cell geometry and the cell packing.

#### Ice propagation as a Markov process

To develop and test our model, we consider the simplest one dimensional tissue array of two cells modelled in [23]. As indicated in Fig 2, there are three possible states, *X*, to be observed in each laboratory experiment: there is no frozen cell, only one cell of two cells is frozen, or both cells are frozen (i.e. *X* = 0, 1, 2 respectively). The transition from state *X* = 0 to *X* = 1, can only be caused by the independent ice formation mechanism *J*^i^, so the rate of loss of unfrozen cells, state *X* = 0, is equal to the rate of growth in state *X* = 1. The transition from state *X* = 1 to *X* = 2 can result from either independent intercellular mechanism *J*^i^ or ice propagation mechanism from the neighbour cell *J*^p^, so the decrease in state *X* = 1 is equal to increase of state *X* = 2 formation plus increase of state *X* = 0. Therefore, we have

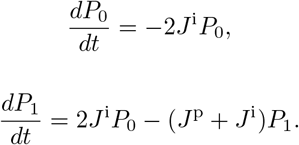

Considering the ice propagation rate *α* = *J*^p^/*J*^i^ and the dimensionless time *τ*, with *P* = (*P*_0_, *P*_1_)^*T*^, we have

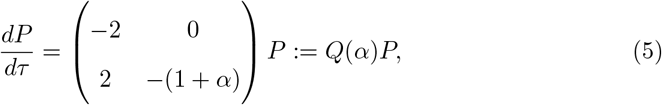

where *Q*(*α*) is the state transition matrix associated with the Markov process. Solving Eq (5) and keeping in mind that *P*_2_ = 1 − *P*_0_ − *P*_1_, we can plot probability as a function of *τ*, as shown in Fig 2. Note that the probability *P*_0_ is independent of *α*.

**Fig 2.**
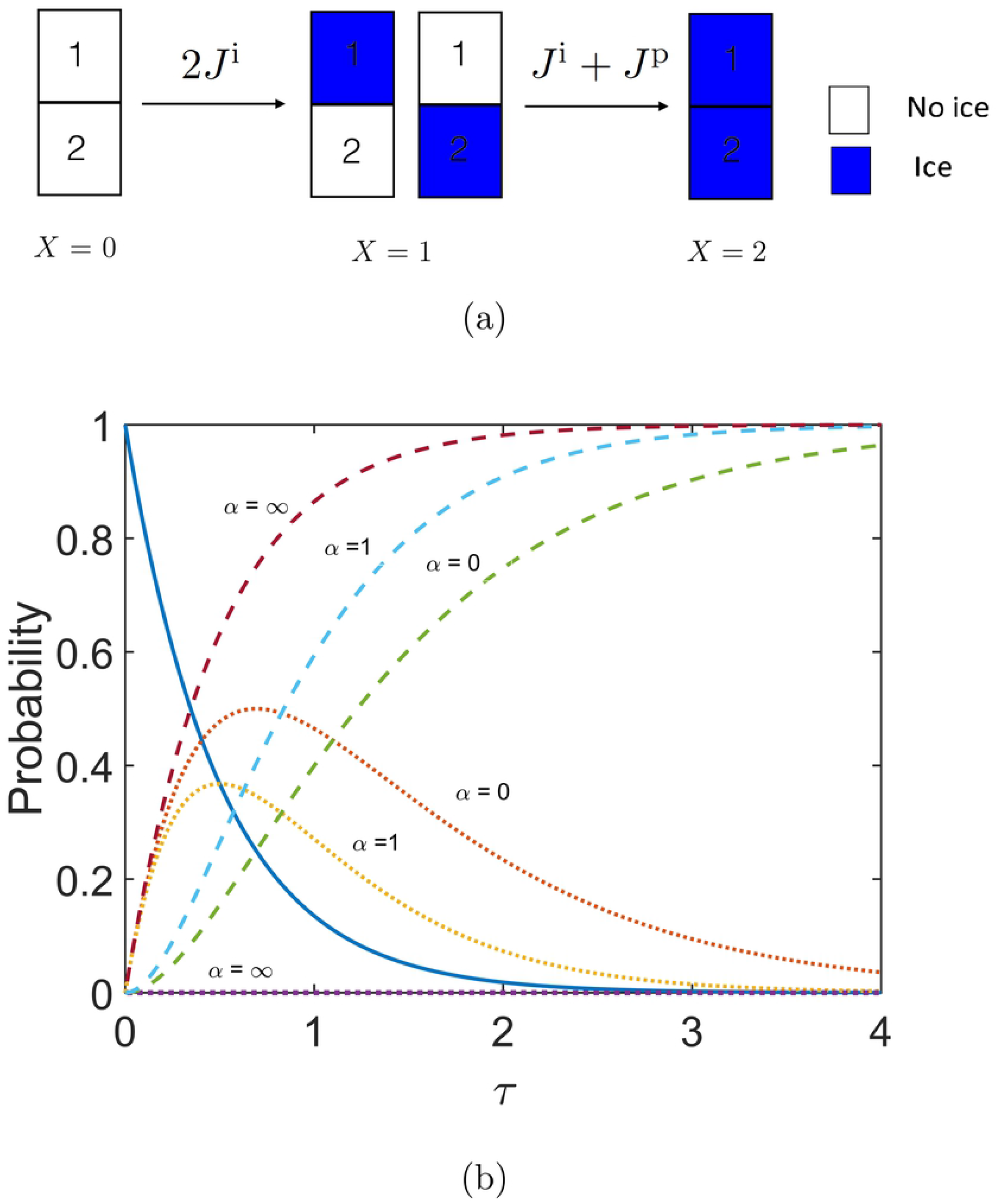
States and their probabilities in two cell tissues. (a) Three possible states, including symmetry, for a two cell tissue with probabilities *P*_0_, *P*_1_, *P*_2_. In particular: state *X* = 0, when there is no ice, state *X* = 1, when either cell number 1 or 2 are frozen (blue), state *X* = 2, when two cells are frozen. Note that rates shown above the arrows account for symmetries. For example, to go from state *X* = 0 to *X* = 1, there are two possibilities for interaction independent ice to form, indicated by the rate 2*J*^i^. To go to state *X* = 2, there is one way to have interaction independent ice and one way to have propagated ice, thus the rate *J*^i^ + *J*^p^. (b) The probabilities of states *X* = 0 (solid line), *X* = 1 (dotted line) and *X* = 2 (dashed line), respectively as a function of the time-like variable *τ* with three rates of propagation, *α* = 0, 1, ∞.

Extending the results from Irimia and Karlsson, now let us consider a new and more complex 2 × 2 cell structure. The possible states and their related rates are illustrated in Fig 3, accounting for symmetries. In this case, for each cell *j*, the maximum number of nearest neighbours is three, including the diagonal neighbour. To determine the probabilities of each state as a function of time in this geometry, an analytical method such as a Markov-chain model is used. Assume that the state *X* ∈ {0,…, *M*} is associated with probability *p_X_*(*t*), where *M* is the maximum number of possible states of observation at a given time. The IIF state *X* includes the frozen and unfrozen individual cells in identical tissue constructs. Since the state space {0,…, *M*} is finite, the transition probability distribution can be indicated by a transition matrix. In the case of *M* = 5, this becomes

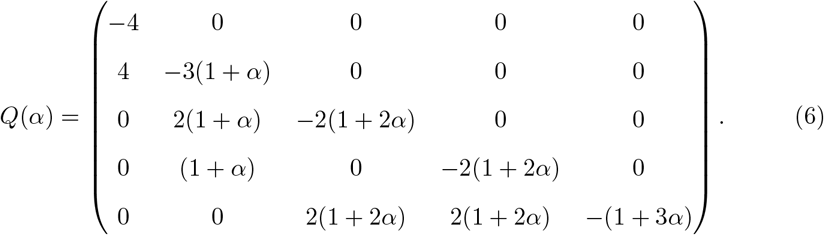

Therefore, if *P* = {*p_X_*}_*X*=0,…*M*_ presents a vector of all probabilities, then the probability of each state can be obtained by solving a Kolmogorov differential equation,

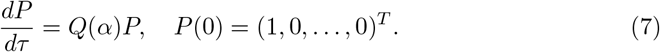

With the above assumptions, the freezing of a tissue construct with *N* cells can be described by a discrete-state, continuous-time, homogeneous Markov chain. The IIF states *X* of the tissue construct comprise all distinct combinations of states (frozen or unfrozen) of the individual cells in the tissue. For example, all possible states and state transitions for a four-cell array are shown in Fig 3 in the form of a directed graph. In an ensemble of identical tissue constructs, the elements of the state probability vector *P* represent the probabilities *p_X_* of each IIF state *X* ∈ {0,…, *M*} at a given time. However, due to computational complexity, the size of system (7) increases exponentially with tissue size and quickly becomes intractable, so a different approach must be used.

**Fig 3.**
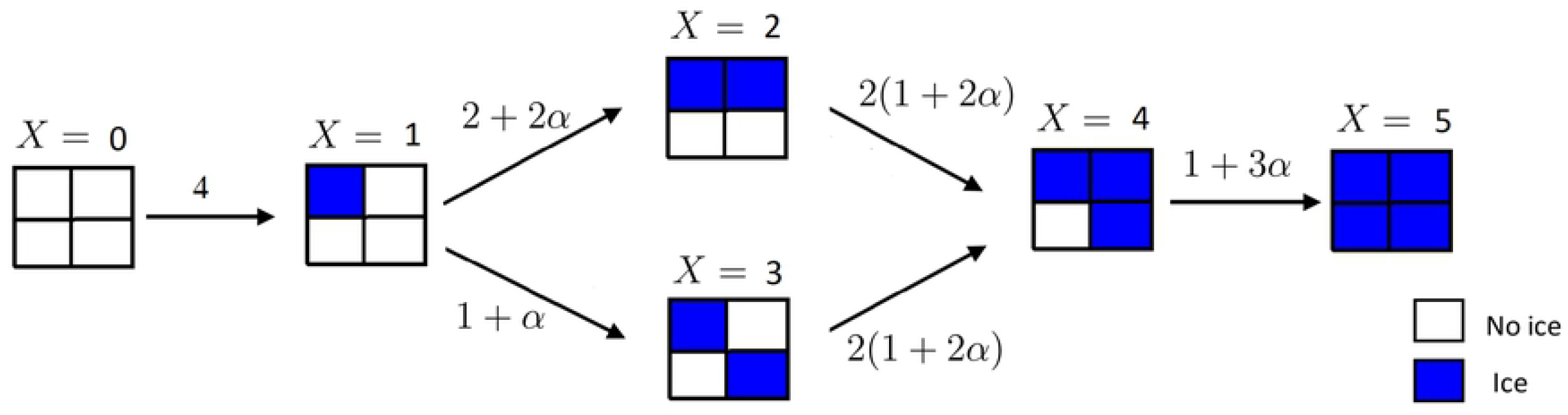
All possible states for 2 × 2 construct with *X* = 0,…, 5 indicated above each state. As in Fig 2, each state depends on the previous state. Symmetries are not shown, but are indicated by the nondimensional rates given above the arrows. For example, there are four symmetric singlet states, so the rate to go from state *X* = 0 to state *X* = 1 is 4. There is only one way to go from a singlet state (*X* = 1) to the opposite corner singlet state (*X* = 3): either via the independent mechanism at nondimensionalized rate 1, or via the propagative mechanism at rate *α*. In contrast there are two symmetrically equivalent ways to arrive at state 2, each with an independent rate of 1 and a propagative rate *α*, totaling 2 + 2*α*, etc.

#### Monte-Carlo Model

The alternative to a Markov chain model is the Monte-Carlo method. Monte-Carlo methods are used to approximate the probabilities of outcomes based on repeated random sampling. For the present model, the probability that an unfrozen cell *j* ∈ {1,…, *N*} may freeze within a time interval Δ*t* is defined by the sum of the probabilities of spontaneous and propagation mechanisms. Following Irimia and Karlsson [24], using Eq (2), Eq (3) and Eq (4) we have

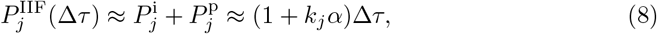

where a sufficiently small Δ*τ* must be chosen to ensure at most one change of state (IIF event) per time-step. Irimia and Karlsson suggest

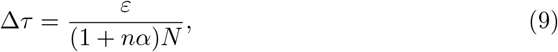

where *n* is the maximum number of nearest neighbours, and *ϵ* is a small probability bound for the IIF probability in the tissue during time interval Δ*τ* [24].

To run numerical experiments, we implemented a Monte-Carlo algorithm as follows. The state of cell *j* is described by

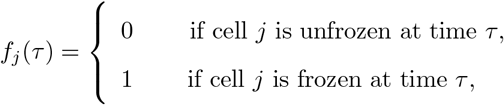

where *f_j_* is a property of the *j*th cell-agent. For every unfrozen cell at each time step Δ*τ*, a random number *r_j_* is generated from the uniform distribution on the interval (0, 1). If 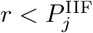 the cell *j* considered as frozen. However, accurate results require a high number of iterations or experiments. This makes the Monte-Carlo method computationally expensive.

To overcome this problem Karlsson and Irimia [24] propose using the Gillespie algorithm [46, 47], which estimates both the reaction (conversion to IIF) location and the time of reaction by two random variables *r*_1_, *r*_2_ compared with the uniform distribution on (0, 1). This means that for a simulation of *N* cells, only *N* time steps are needed, a considerable savings in computational time.

To implement this algorithm, assume that the overall rate at time step *i* is as follows

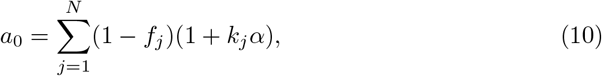

where *N* is the number of cells in the tissue. Then, the next time step is

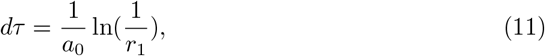

and the specific cell is to be frozen at the time *τ* + *dτ* is given by the *j* that satisfies 260

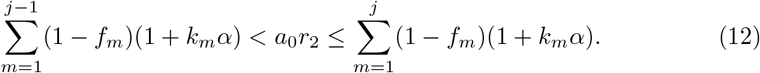

We implemented both the standard Monte Carlo and Gillespie algorithms within the construct of PhysiCell, and note that when *N* < 100, their computational cost is similar.

To validate our implementation of the Monte-Carlo methods, we calculated the probability of each of the six states as a function of time for the 2 × 2 structure (Fig. 3), shown in Fig. 4, where we compare exact Markov and numerical Monte-Carlo models. Using parameters from Irimia and Karlsson (e.g. *α* = 10.4), we compared mean results from 10^5^ Monte Carlo iterations with 1744 total time steps. As the number of iterations or repeated experiments increases, the accuracy of the Monte-Carlo method improves and approaches the theoretical solution from Markov algorithm Fig 5.

**Fig 4.**
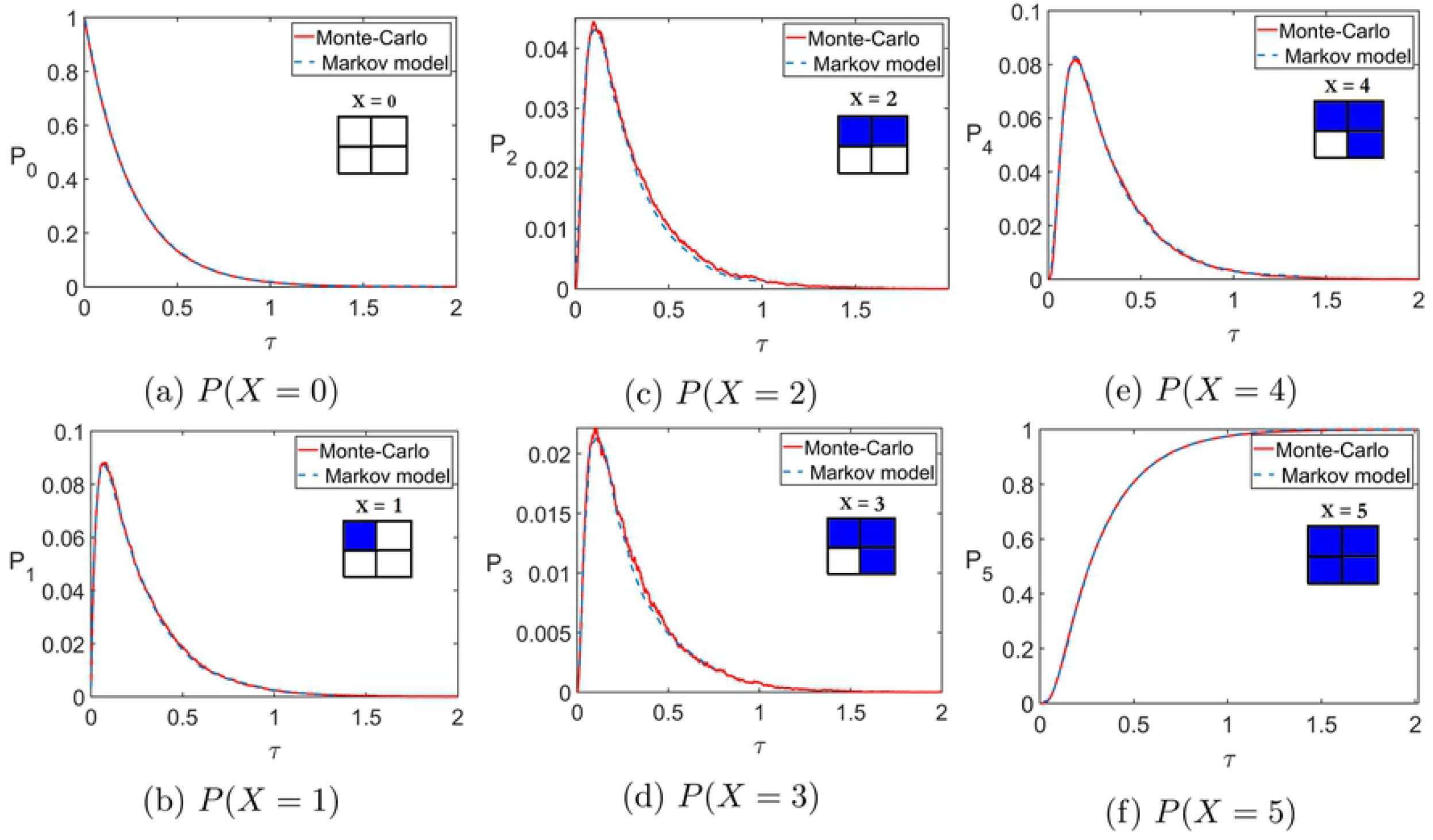
Exact (Markov chain) and numerical (Monte-Carlo) predictions of the probabilities of each state in the 2 × 2 structure (c.f. Fig 3), with propagation rate *α* = 10.4, implemented within the PhysiCell agent-based code. Inset is the associated diagram for each state, with blue indicating ice.

**Fig 5.**
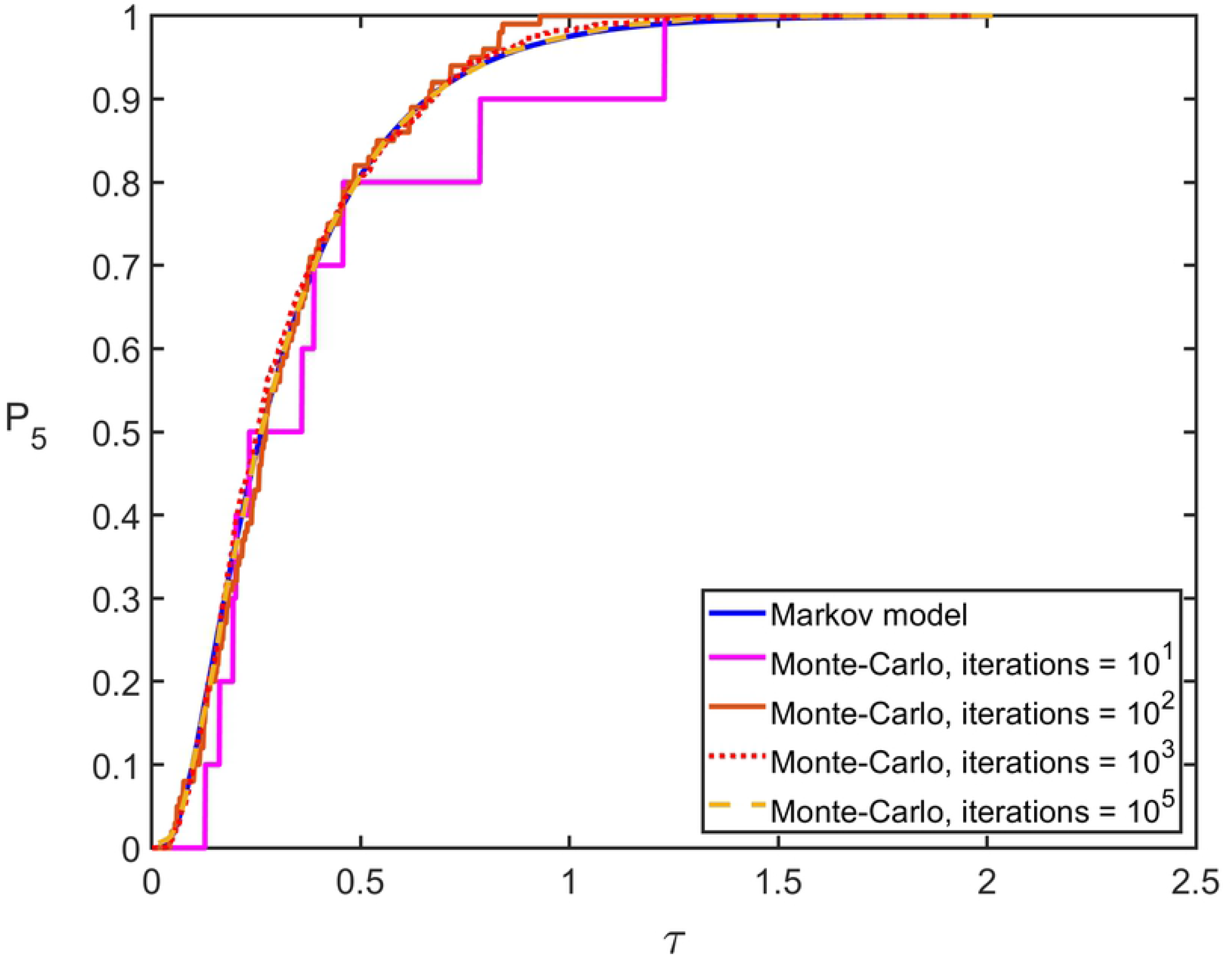
Exact (Markov chain) and numerical (Monte-Carlo) predictions of the probability of state *X* = 5 in the 2 × 2 structure (c.f. Fig 3), with propagation rate *α* = 10.4 and different numbers of iterations.

## Water Transport model

Cell water content plays a critical role in survival during freezing [12]. At slow cooling rates, extreme dehydration and extended exposures to high osmolality solutions are associated with deleterious “solution effects” [48], while at high cooling rates the damage of ice formation is considered the dominant factor [15]. This is because, at high cooling rates, water has insufficient time to leave the cell, causing the intracellular milieux to be supercooled; and, as temperature decreases, the probability of ice formation inside the cell increases along with the probability of cell damage. From these two modes of damage, an optimal cooling rate can be summarized as “fast enough to minimize solution effects, but not so fast as to cause ice formation.” This present work focuses on understanding ice formation, and as *J*^i^ is the mechanism of ice formation from inside the cell, it is a function of water content of the cell. So, a model that describes water transport during freezing is needed.

We consider a nonequilibrium water transport model introduced by Mazur [12]. Transport across the cell membrane is driven by chemical potential differences between the intracellular and extracellular solutions [49]. Hence, the kinetics of water loss at subzero temperatures is:

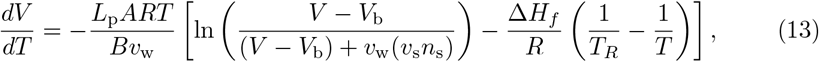

where *V* is cell volume; *R*, the universal gas constant; *T*, absolute temperature; *A*, surface area of the cell; *B*, the cooling rate; *v*_w_, partial molar volume of water; *V*_b_, osmotically inactive cell volume; *v*_s_ = 2, the dissociation constant for NaCl; *n*_s_ number of moles of salt in the cell; Δ*H_f_* the latent heat of fusion of water; *T_R_* the equilibrium freezing temperature for pure water 273.15 K; and *L*_p_, the hydraulic permeability. The temperature dependence of *L*_p_ is given by an Arrhenius model with

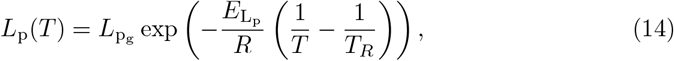

*L*_P_g__ is the permeability of water at *T_R_*, and *E*_L_p__ is the activation energy for the water transport process. For commonly implemented constant-rate cooling protocols, the temperature is given by

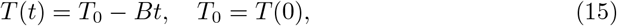

where the transformation from temperature to time can be implemented using the cooling rate *B*.

### Non-dimensional time

Up to now, our model was defined and evaluated in terms of the time-like variable *τ*, but in order to compare the introduced model with experimental results, we need to obtain *τ* as a function of time or temperature. From Eq (3), *τ* can be evaluated if the cell surface area A and the independent rate *J*^i^(*T*) are known. Karlsson *et al*. [17, 20, 21, 50] define

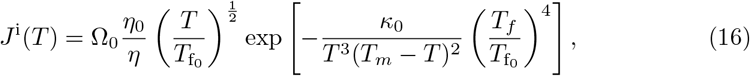

where the subscript ‘0’ refers to isotonic conditions, Ω_0_, *κ*_0_ are kinetic and thermodynamic coefficients respectively,

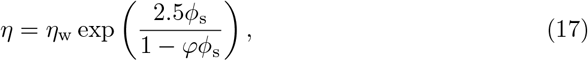

is the cytoplasmic viscosity as a function of the volume fraction of salt, *ϕ_s_*, *φ* = 0.609375 is the interaction parameter, and *η*_w_ is the temperature dependent viscosity of pure water [20],

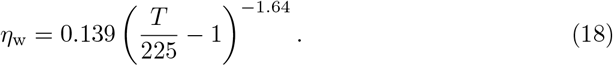

The equilibrium freezing temperature of the cell cytoplasm [19]

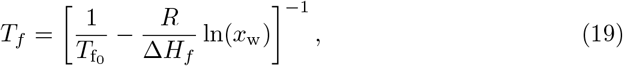

is a function of the mole fraction of water,

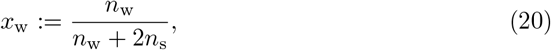

and where *n*_w_ and *n*_s_ are moles of intracellular water and dissociated salt, respectively. Note that it is this mole fraction that requires the water content of the cell, modelled above using Eqs (13) to (15).

### Ice expansion and tissue remodeling

As a first approximation for capturing the potential disruption of tissues due to the sequential solidification of the intracellular spaces, we used the following approach. First, we modeled the propagation of ice in the tissue structure as described above. Then for any particular Monte-Carlo simulation, we then made a list of times *τ_j_* at which cell *j* solidified. Then we converted to real time, *t_j_*. Then we used model Eq (13) evaluated at time *t_j_* to determine the intracellular water content at the time that the cell solidified. This water was assumed to turn entirely into ice with a volume expansion of 1.09. Then running in real, dimensional time, we used the PhysiCell mechanical force model with default parameters for our tissue construct [40], except where the volume state of the *j*th cell was changed at time t*j* to have its new, expanded volume. This process was repeated for all *j* ∈ *N*, and the force model was allowed to run as usual throughout the entire freezing experiment. For simplicity of presentation (i.e. it is difficult to visualize the mechanical disruption in 3D tissue constructs), we only performed this procedure on our 22 cell hepatocyte monolayer disks.

## Results and Discussion

### Nondimensional time validation in single cells

While the value of the nondimensional time *τ* depends on the exact form of *J*^i^(*t*) and eventually on a given cell type and cooling rate, in a homogenous tissue, the relative sequence and spatial distribution of IIF events depends only on the magnitude of the nondimensional propagation rate *α*. Therefore, here we implement the model to study ice propagation in rat hepatocyte tissue constructs.

The biophysical parameters for hepatocytes are given in Table 1 and were taken from [20, 51]. In Fig 6, the normalized cell volume is plotted as a function of temperature for an average rat hepatocyte. As can be seen from Fig 6, the cell water content depends strongly on the cooling rate. At a moderate cooling rate such as *B* = 50 °C/min, the cell loses almost 50% of its water, while for a rapid cooling rate, *B* = 400 °C/min, the cell retains about 85% of its original water content. Fig 7 illustrates the monotonic (and therefore invertible) relationship between *τ*, time and temperature for rat hepatocytes, respectively. Here, the required parameters for updating the *J*^i^ model (Eq (16)), specifically the water volume of the cell that manifests in Eq (19), are updated from the water transport model (Eq (13)). When comparing or fitting model predictions below (in which kinetics are described in terms of the nondimensional time variable *τ* to experimental data, which describe probabilities of ice as a function of temperature), and the relationship shown in Fig 7 was used to convert between units of temperature and nondimensional time.

**Table 1.** Water transport and IIF parameters of individual rat hepatocytes [20].

| Parameter | Value |
| --- | --- |
| $L_{pg}$ | $15 \times 10^{-13} \text{ m}^3 \text{ N}^{-1} \text{ s}^{-1}$ |
| $E_{Lp}$ | $342 \text{ kJ mol}^{-1}$ |
| Cell Diameter | $21.2 \text{ } \mu\text{m}$ |
| $V_b$ | $0.51$ (unitless) |
| $\Omega_0$ | $110 \times 10^8 \text{ m}^{-2} \text{ s}^{-1}$ |
| $\kappa_0$ | $14 \times 10^8 \text{ K}^5 \text{ s}$ |

**Fig 6.**
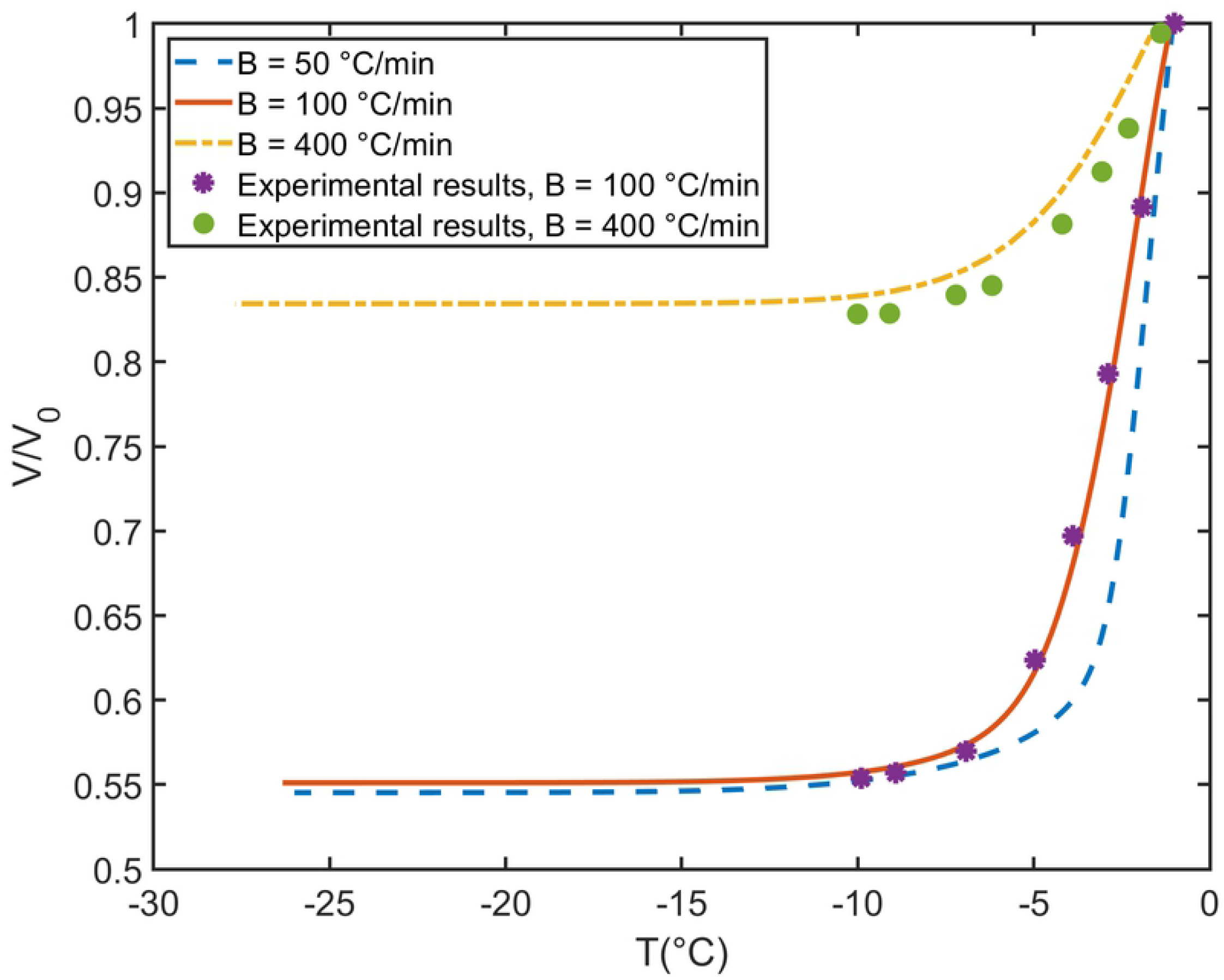
Comparison of theoretical and experimental results for normalized total individual rat hepatocyte volume as a function of temperature at different cooling rates. The experimental results were digitized from [20].

**Fig 7.**
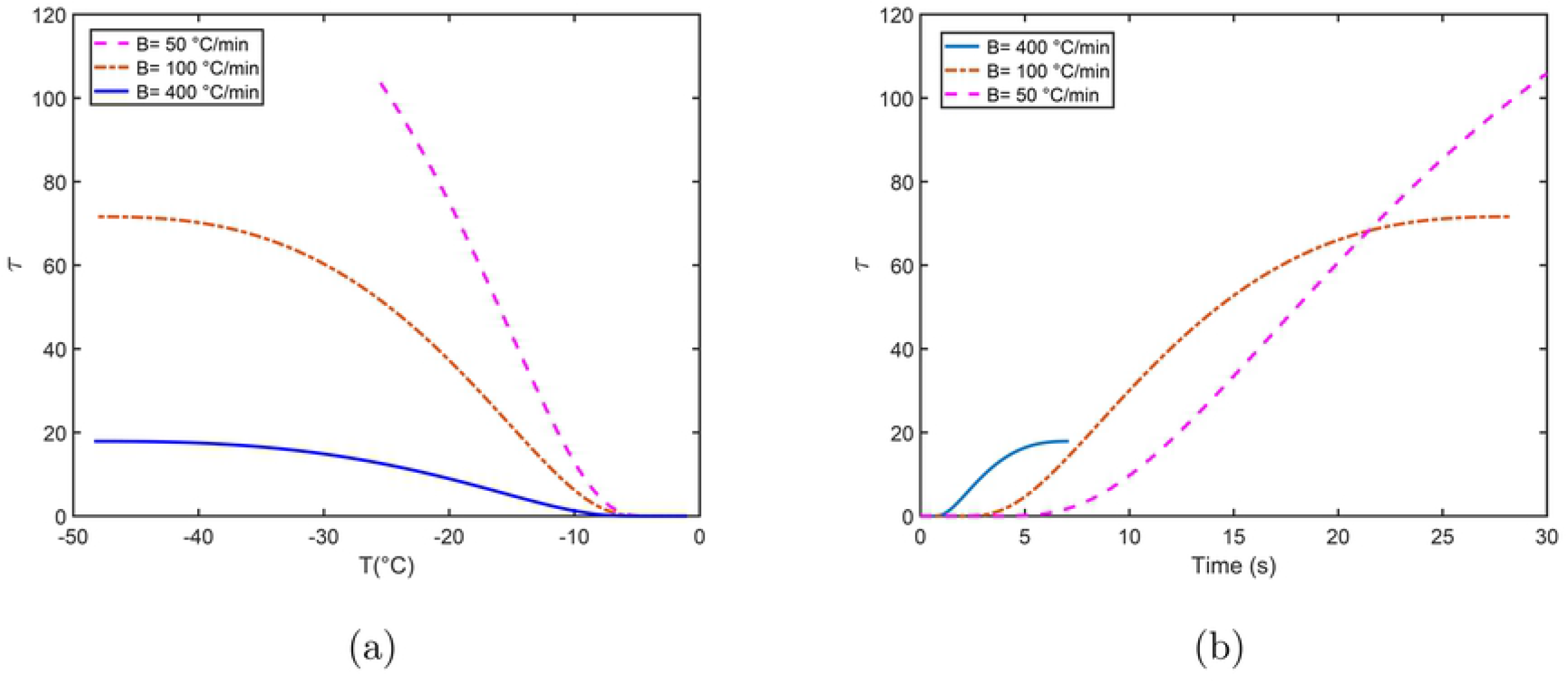
Nondimensional time *τ* for rat hepatocytes as a function of (a) temperature in Celsius and (b) real time in seconds.

Fig 8 shows the comparison between theoretical and experimental results obtained from using a cryomicroscopy system at different cooling rates [51]. The theoretical IIF probability results first were obtained based on dimensionless *τ*, then *τ* was converted to temperature using Fig 7. The small difference between our results and the experimental results, indicate a good approximation using our method.

**Fig 8.**
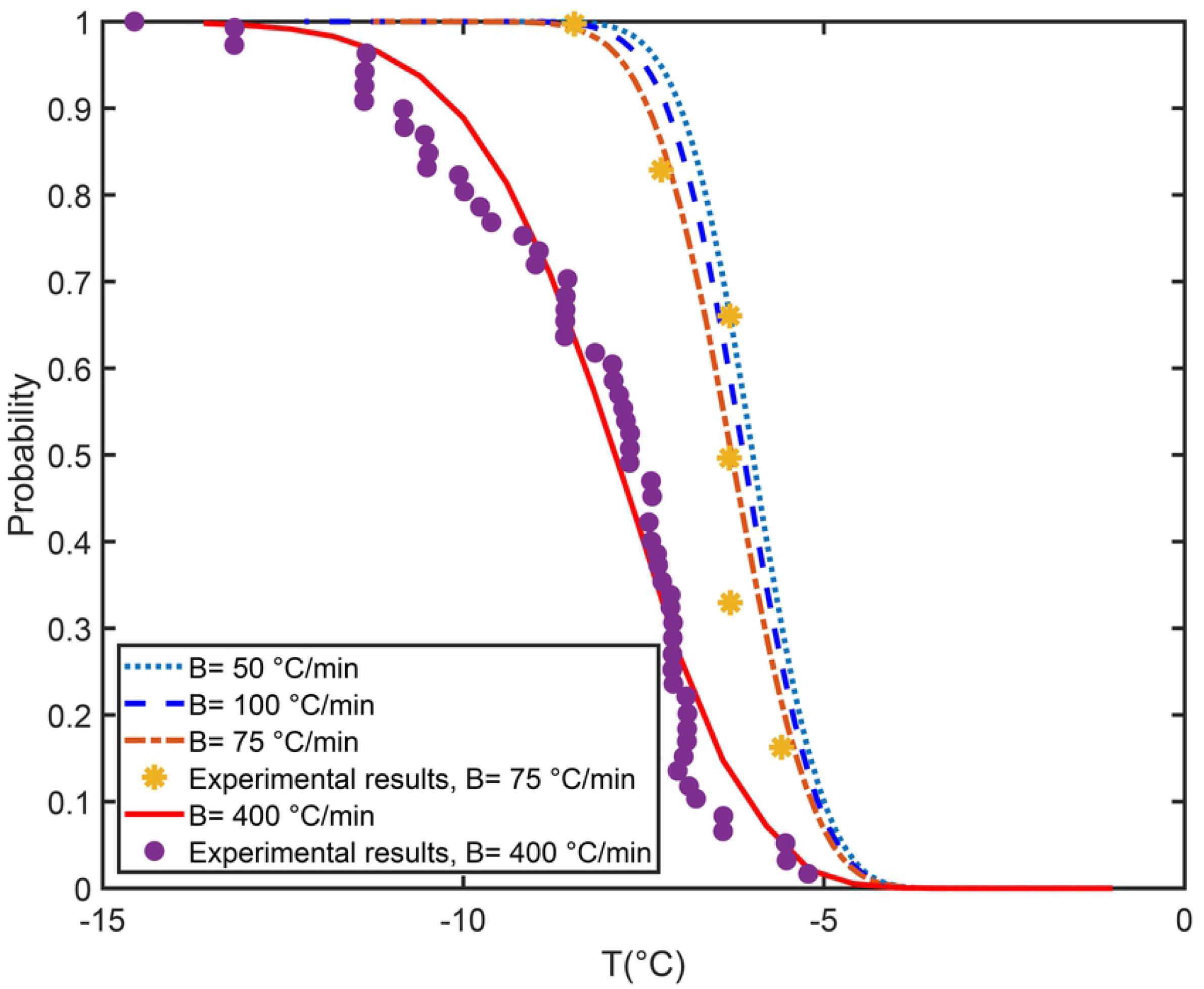
The probability of IIF for isolated rat hepatocytes with theoretical and experimental results and different cooling rates. The experimental results were digitized from Toner *et al*. [20, 51].

### Rat Hepatocyte Disk Monolayers and Spheroids

#### Disk Monolayers

In this example, we first consider a lattice free disk shaped monolayer of rat hepatocyte with 22 cells, Fig 1. Simulations were performed to investigate the effects of nondimensional ice propagation parameter *α*, as well as *R*, the radius of the cell neighbourhood, or *k*, the maximum number of neighbours, on the model. The cell properties are given in Table 1. In Fig 9, two different values of *R* = 21 or *R* = 22 *μ*m with associated *k* = 2 or *k* = 7, respectively, are studied. The results indicate that when *R* and *α* decrease, the intracellular ice propagation, *J*^p^, decreases as well. That means for the two different *R* and the same *α*, the number of cells that freezes from ice propagative IIF is higher for higher *k*. So, *J*^p^ depends on the number of neighbour cells, *k*, and *α*.

**Fig 9.**
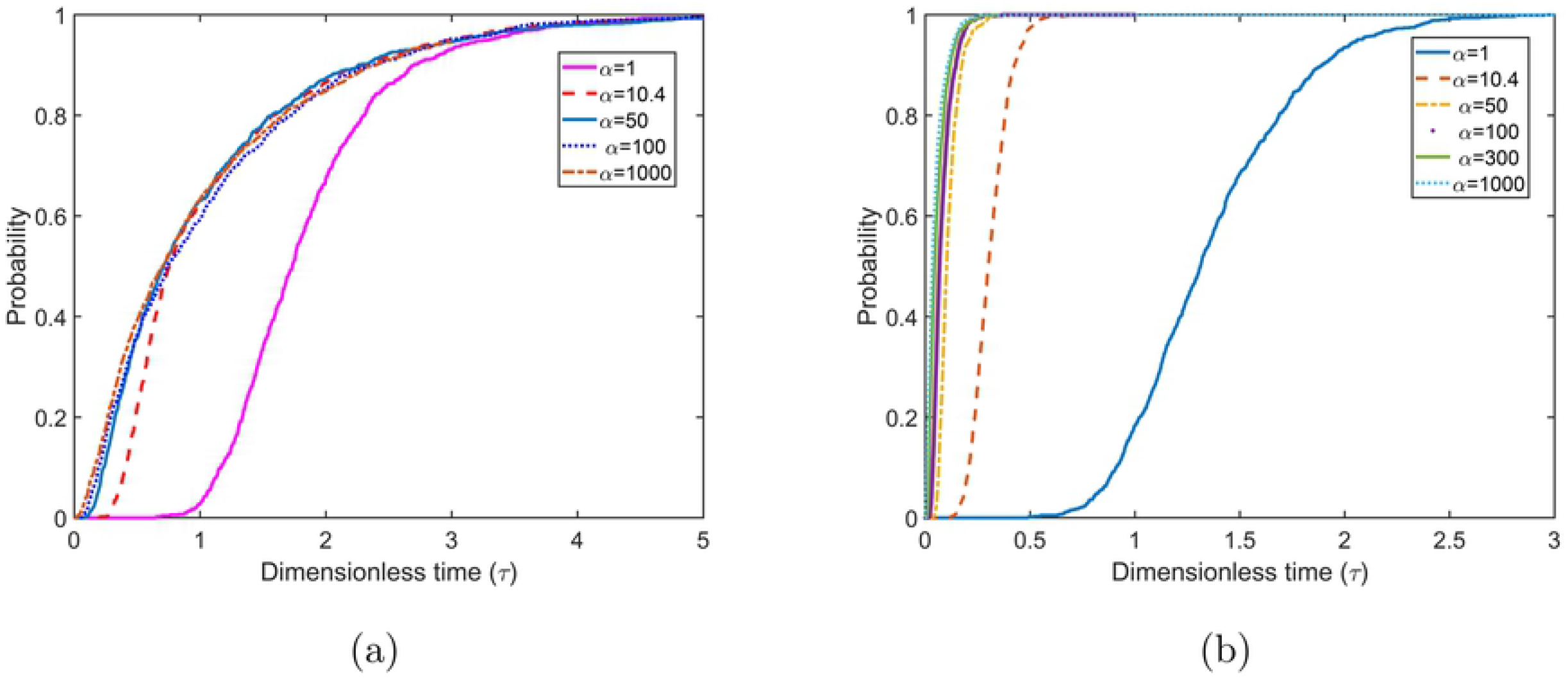
IIF in disk monolayers. (a) Probability of IIF for 22 cell hepatocyte monolayer with (a) *R* =21 *μ*m. (b) *R* = 22 *μ*m. Note that the hepatocyte diameter used is 21 *μ*m.

As can be seen from Fig 9, a smaller *τ* is obtained for higher *R*. This demonstrates that the higher connection between cells induces more ice formation due to ice prorogation and reduces the ice formation due to the intracellular ice nucleation, *J*^i^. For fixed *R* and different α there is a threshold such that increasing *α*, does not effect the solution, and Fig 9 illustrates that the threshold value for both cases is approximately 50. This is due to higher number of cells connection and also higher number of gab junctions that facilitate the ice propagation [52]. However, *J*^i^ is independent of the number of neighbour cells *k* and *α*. In fact, as the cooling rate increases to infinity the intracellular ice nucleation, *J*^i^, decreases to zero:

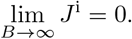

Acker and McGann proposed that intracellular ice propagation is harmless and the damage is only caused by intercellular ice nucleation, *J*^i^ [53, 54]. This suggests that for high cooling rates the IIF damage is reduced by eliminating *J*^i^.

In order to study the ice propagation path, we consider a pre-frozen cell located as indicated in Figs 10 and 11. Two cases were studied: 1) a pre-frozen cell in the centre, 2) a pre-frozen cell in the outside boundary of the monolayer. For each case the Gillespie method was used with 100 iterations and the neighbour list for each cell is defined based on cell to cell membrane contacts. Since, each agent or cell has property of being frozen or unfrozen, the probability that the specific cell freezes and the probability that the ice propagates in a certain path in the monolayers is indicated in Figs 10 and 11. These figures indicate the probability distribution path in the tissue for both cases. We found that when the pre-frozen cell is in the centre, the ice propagates symmetrically towards outside of the monolayer. Whereas, when the pre-frozen cell was on the border, the ice is more likely to propagate to the centre of the monolayer. These results align with the intuition that ice propagates through highly networked cells, and thus does not propagate quickly along the outer surface of the tissue structure.

**Fig 10.**
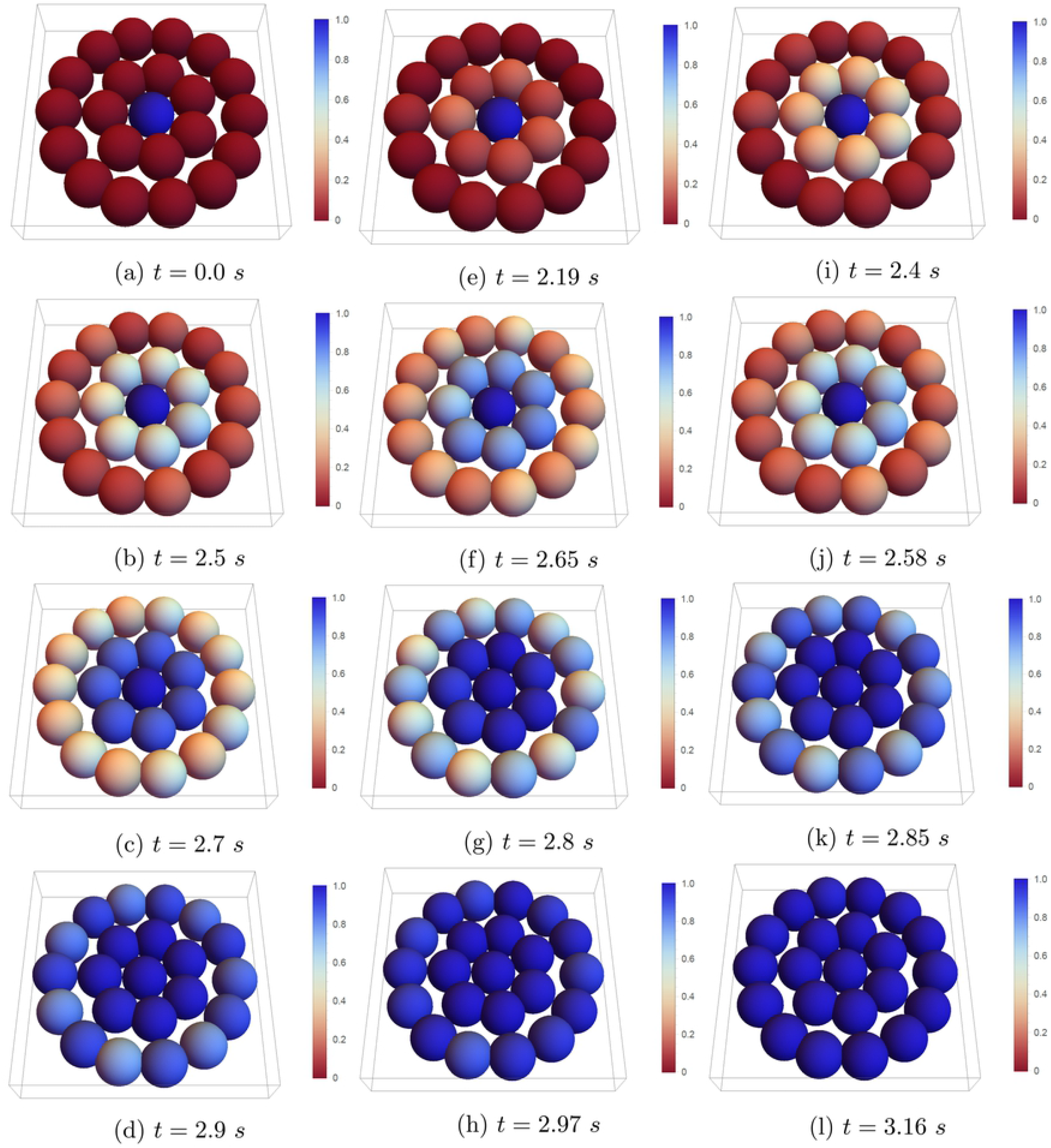
Probability distribution of ice propagation in different time steps of rat hepatocyte monolayer with 22 cells and a pre-frozen cell in the center with *α* = 10.4 and *B* = 100 °C/min.

**Fig 11.**
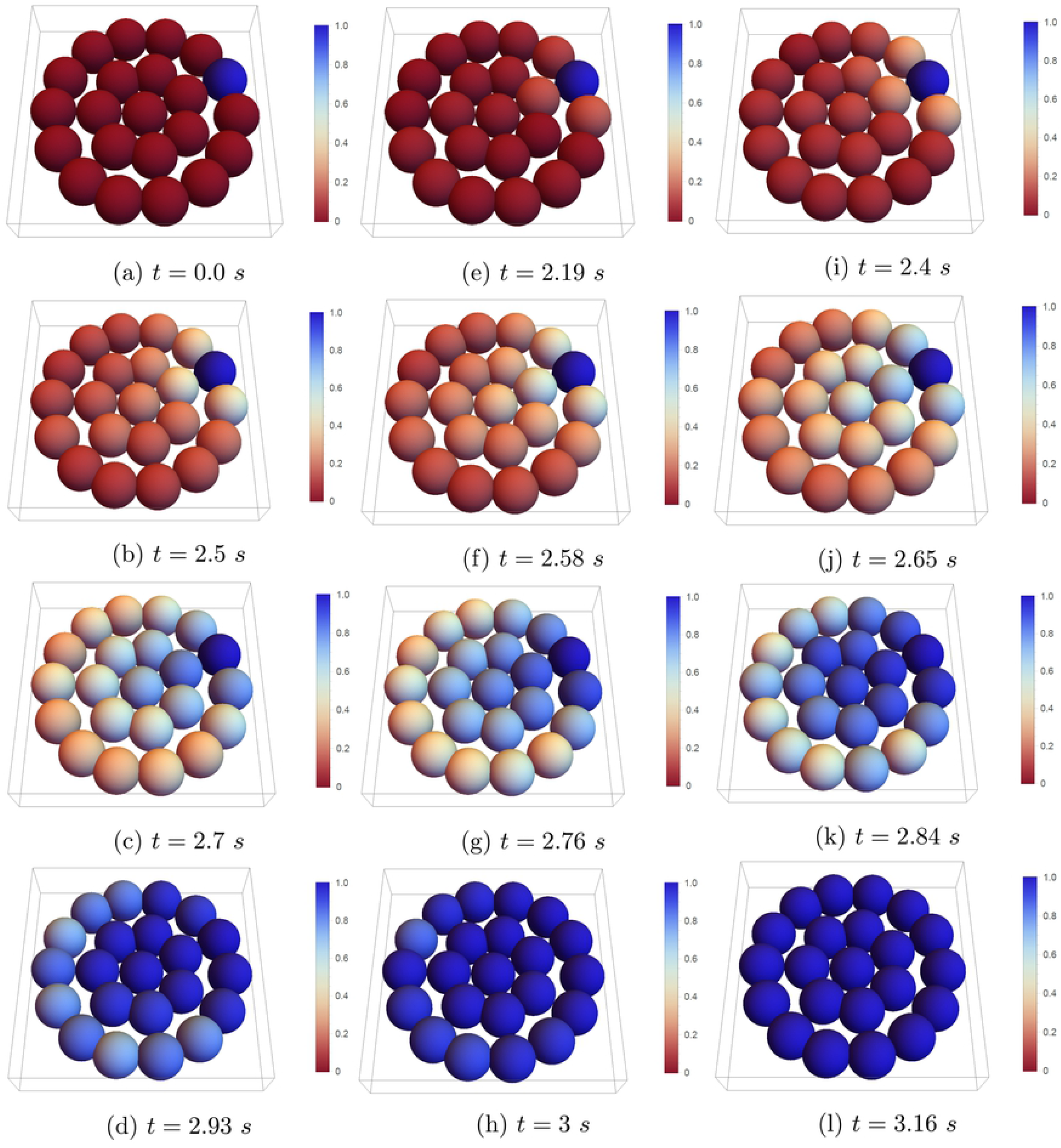
Probability distribution of ice propagation in different time steps of rat hepatocyte monolayer with 22 cells and a pre-frozen cell in the corner with *α* = 10.4 and *B* = 100 °C/min.

In order to study the mechanical disruption of volume expansion we performed a small set of numerical experiments on a 2D tissue structure to enable better visualization. In these experiments the sudden expansion of intracellular water into ice was modelled using PhysiCell’s built in mechanics models experimentally validated for liver structures [55]. In our simple demonstration, shown in Fig 12, the volume change during the dehydration under moderate and fast cooling rates. Upon the solidification, two things change. First the cell ceases to lose more volume, and second any intracellular water is converted to ice at the expansion of 1.09 [56]. This cessation of volume loss is apparent in the volume of in the solidified “first” (top right corner) cell in Fig 12 vs one of the later cells to solidify, where the 10-50% of a volume differential is apparent. In our presented results, it is clear that there are two different ice propagation mechanisms at work at the moderate and fast cooing rates. For *B* = 50 °C/min, ice propagates sequentially from cell to cell, creating a region of non-reactive cells that do not continue to shrink. This causes potential fissure points across the plane of ice propagation. In contrast, at the fast cooling rates (*B* = 400 °C/min) there is less order in the position of cells that solidify, and there is also reduced volume loss due to the shorter time course. This is in line with expectations that cells that are supercooled due to insufficient water loss are more likely to form heterogeneous intracellular ice independent of any propagative mechanism. From an intercellular mechanics point of view, this fast cooling rate manifests in more individual cells separating from their neighbours, suggesting that at higher cooling rates, a different mechanism of ice damage may occur. Note however, as we observe here and below, ice propagates preferentially through well-networked cells. This suggests that there may be cases where some tissue structures, or even some specific cell types, that are likely to get ice and cause mechanical disruption to that portion of the tissue. Moreover, there may be more realistic tissue structures that are more sensitive to these disruptions. Finally, our approach is a first approximation of the actual mechanical dynamics of solidification in tissues. One expects, for example, temperature and state dependence of intercellular force mechanics, as well as the dependence on the state and domain of extracellular ice. If some of the extracellular spaces is already solid, cells may be bound in complicated ways to each other and already solidified portions. More research and careful experimentation is needed on this aspect.

**Fig 12.**
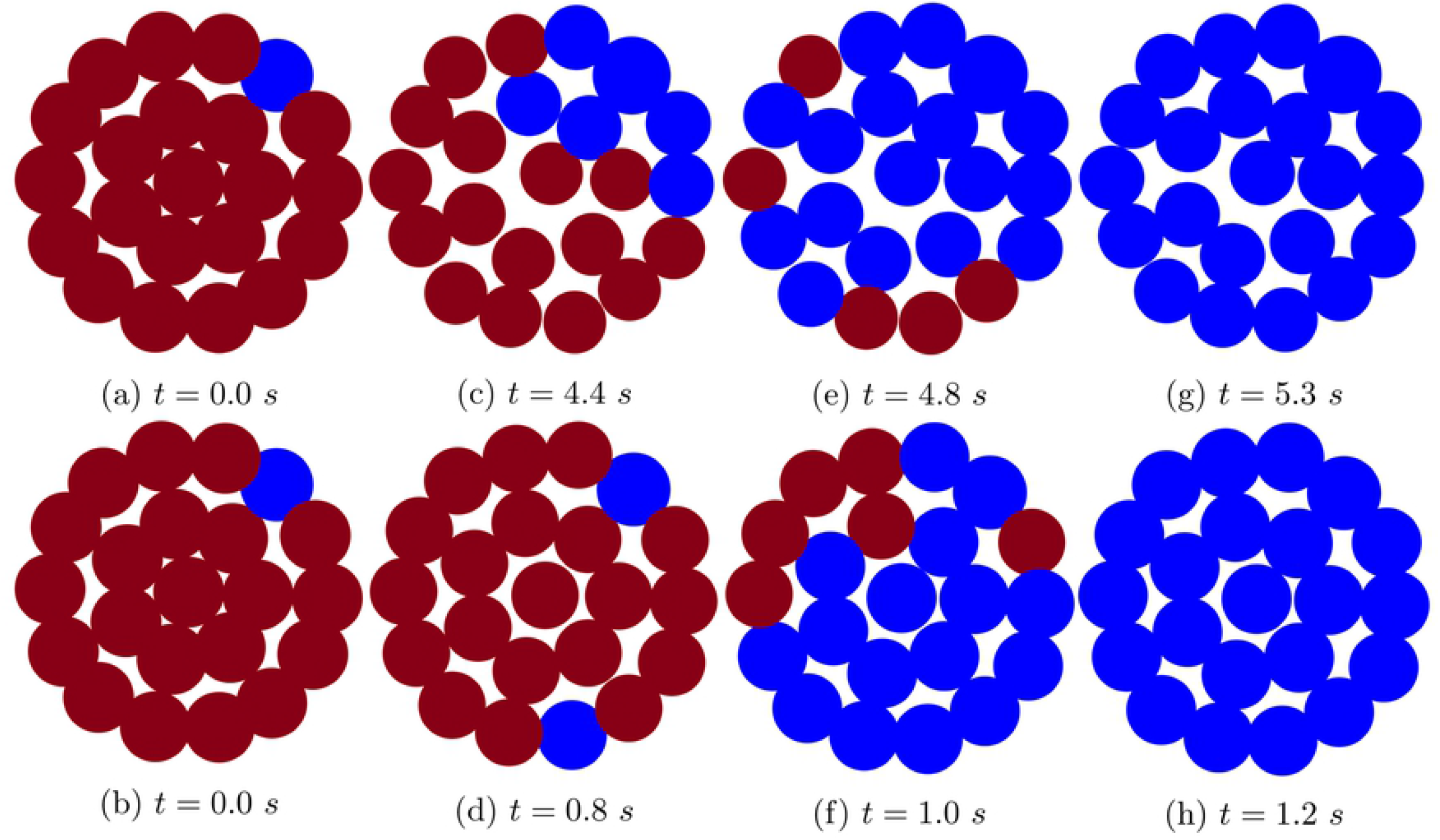
Deformation of rat hepatocyte monolayer with 22 cells at four indicated time points with *α* = 10.4 under two cooling rates, *B* = 50 °C/min (top) and *B* = 400 °C/min (bottom). Red and blue colors indicate unfrozen and frozen cells, respectively.

#### Hepatocyte Spheroids

Rat hepatocyte spheroids are generally 100-200 *μ*m in diameter, are roughly spherical, and contain approximately 100 hepatocytes [57]. Here we emulated this in our model by creating a 3D lattice-free rat hepatocyte spheroid of approximately 100 *μ*m in diameter with 103 cells and started our simulations with a pre-frozen cell on the border of the spheroid. The probability of ice propagation in three different layers of the spheroid is illustrated in Fig 13 at different time steps. In Fig 13, *a′* indicates the whole spheroid with 103 cells, while in *b′* and *c′* one and two layer from the top of the spheroid have been removed respectively, in order to present more views of the inside the spheroid. In this case, analogous to the monolayer case, ice propagates towards the centre of the spheroid where there are more connections between cells.

**Fig 13.**
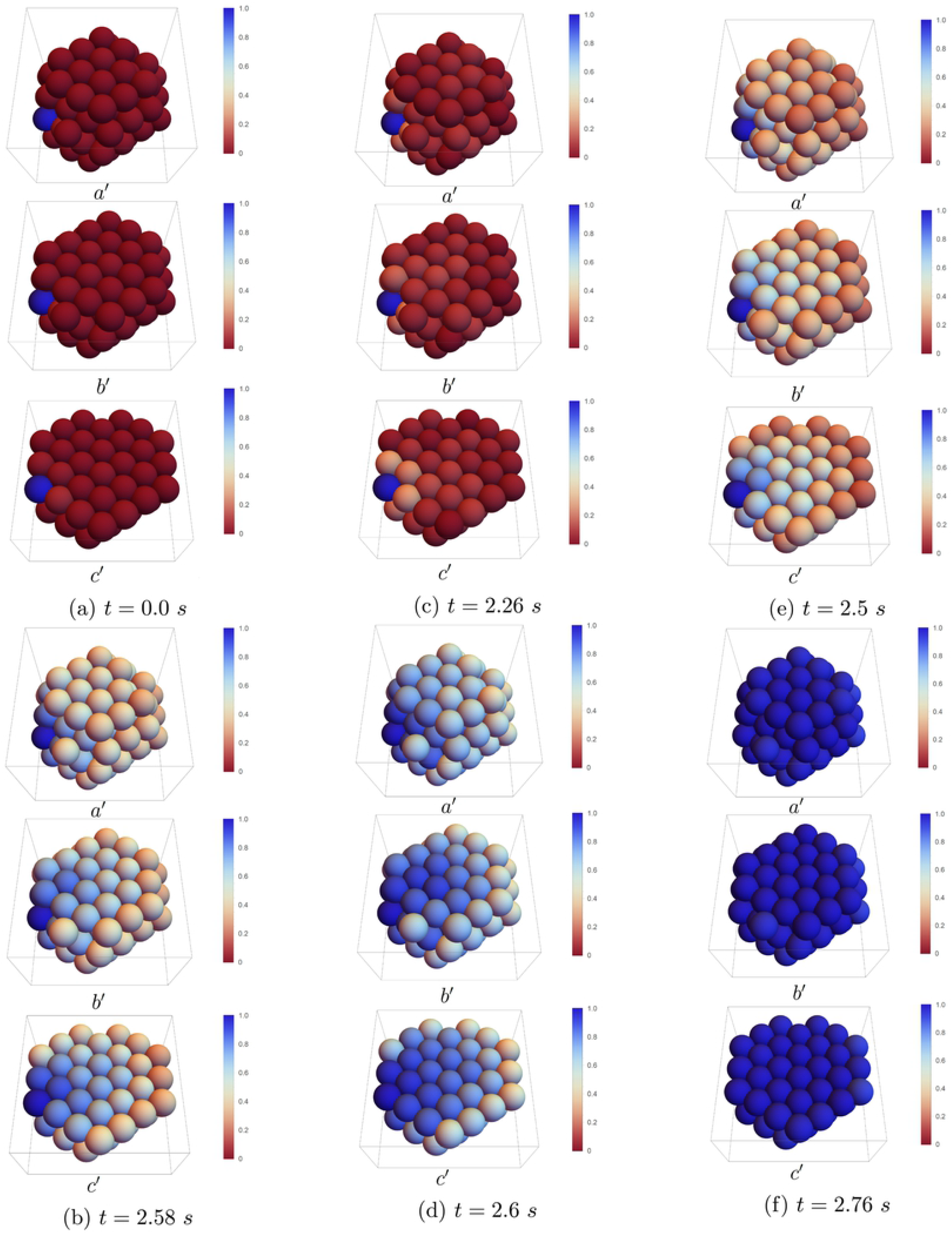
Probability distribution of ice propagation in different time steps of rat hepatocyte spheroids with 103 cells and a pre-frozen cell in the corner of the spheroid with *α* = 10.4 and *B* = 100 °C/min. The whole spheroid is indicated in *a′*, one layer has been removed in *b′*, and two layers have been removed in *c′*.

### Hepatocyte tissue slab

Next, we consider a larger three dimensional homogenous tissue slab comprised of rat hepatocytes with a lattice based cell arrangement of 31 × 11 × 7 cells (approximately 651 *μ*m × 231 *μ*m × 147 *μ*m) with one initial frozen cell seeded in one of three locations: in a corner, in the centre of the top face of the tissue, and in the centre of the tissue. For each case the Gillespie method was applied over 100 iterations. Figs 14–15 indicate the probability of ice as a function of location in the tissue. In this construction, the maximum number of neighbours for each cell is *k* = 8, which becomes reduced if the cell is on a face, edge, or corner of the tissue slab.

**Fig 14.**
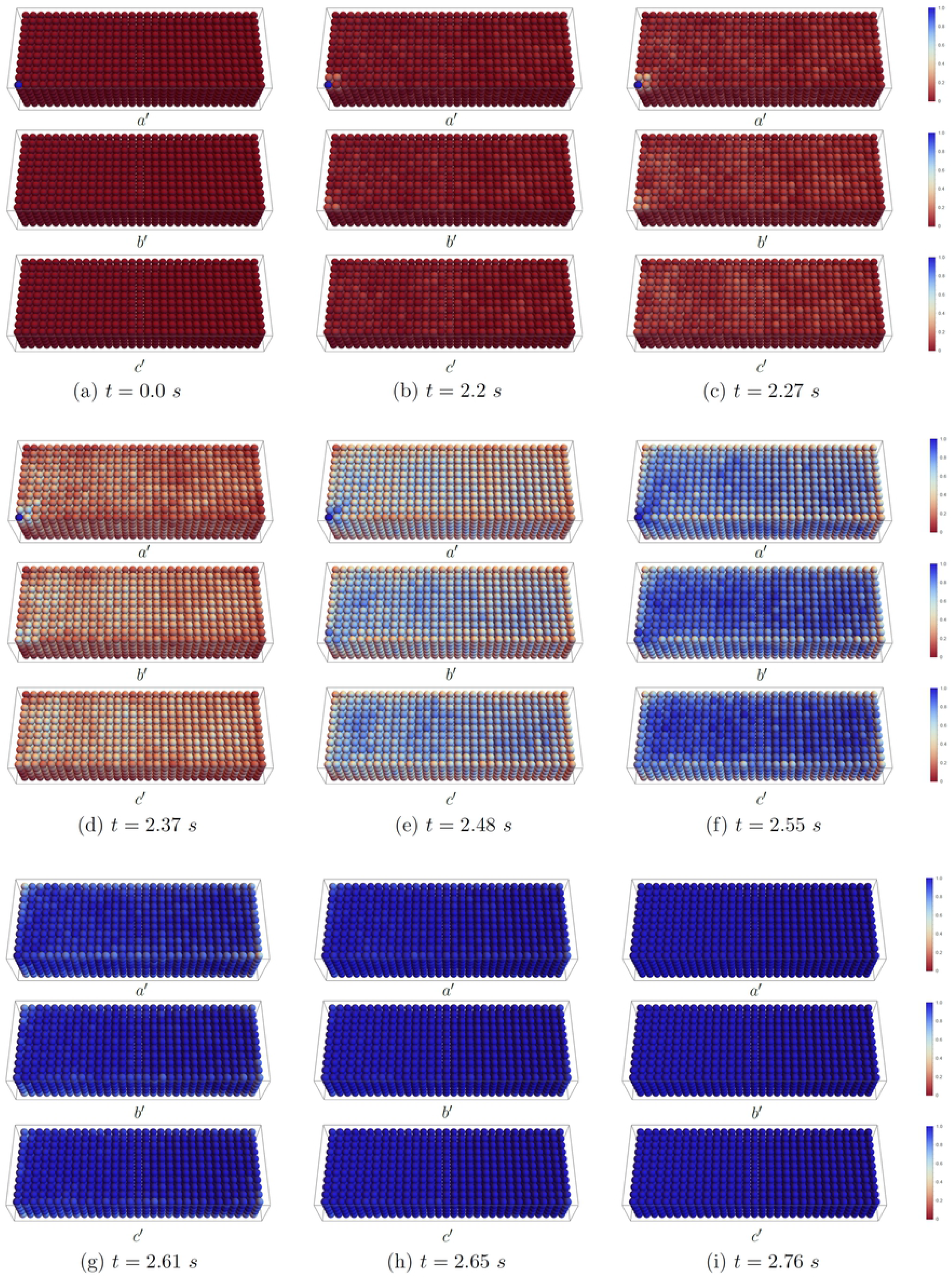
Probability distribution of ice propagation in different time steps of a large tissue with 2387 cells and a pre-frozen cell in the corner of the tissue with *α* = 10.4 and *B* = 100 °C/min. The whole tissue is indicated in *a′*, one layer has been removed in *b′*, and three layers have been removed in *c′*.

**Fig 15.**
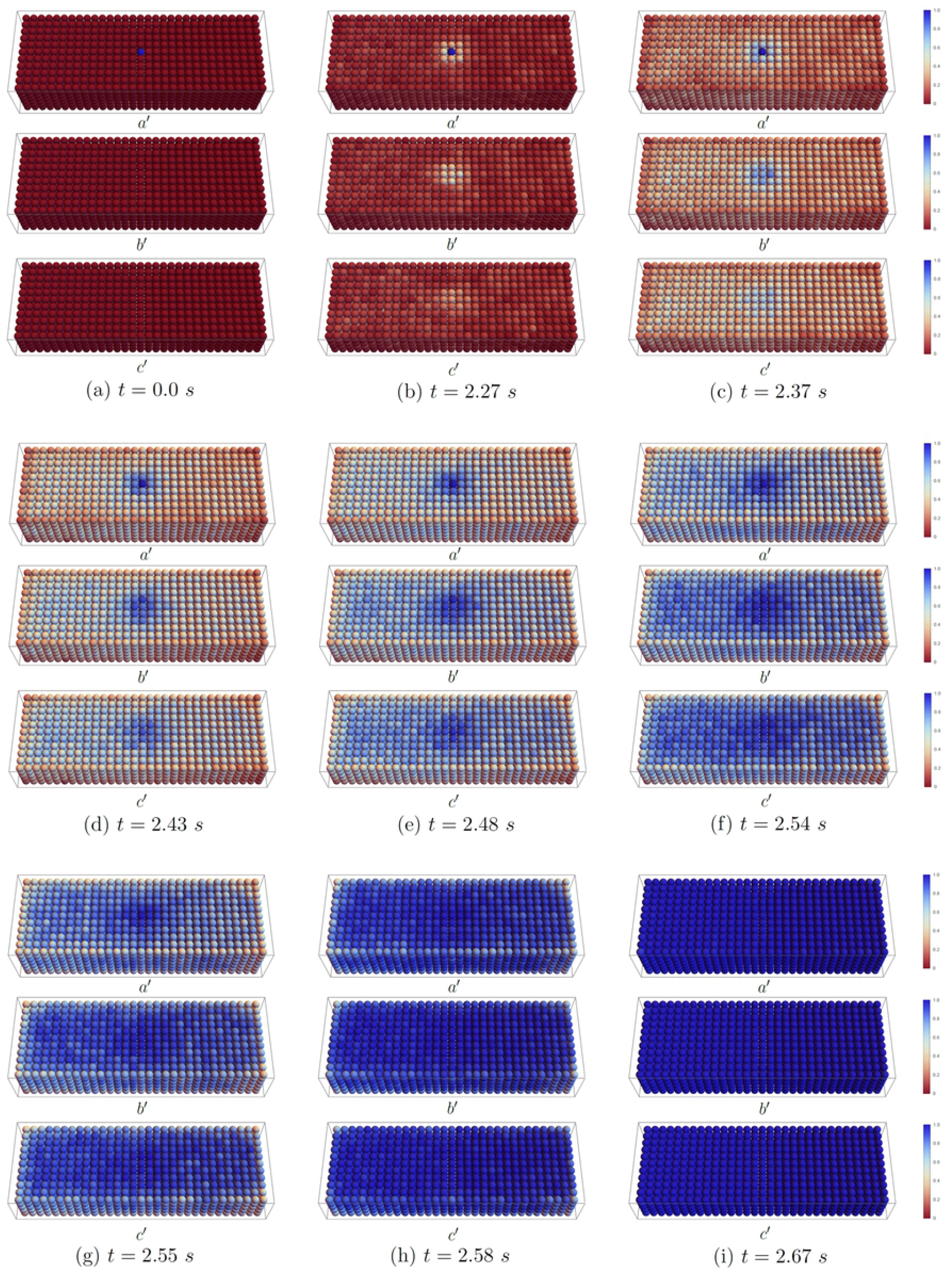
Probability distribution of ice propagation in different time steps of a large tissue with 2387 cells and a face-seeded cell on the top of of the tissue with *α* = 10.4 and *B* = 100 °C/min. The whole tissue is indicated in *a′*, one layer has been removed in *b′*, and two layers have been removed in *c′*.

**Fig 16.**
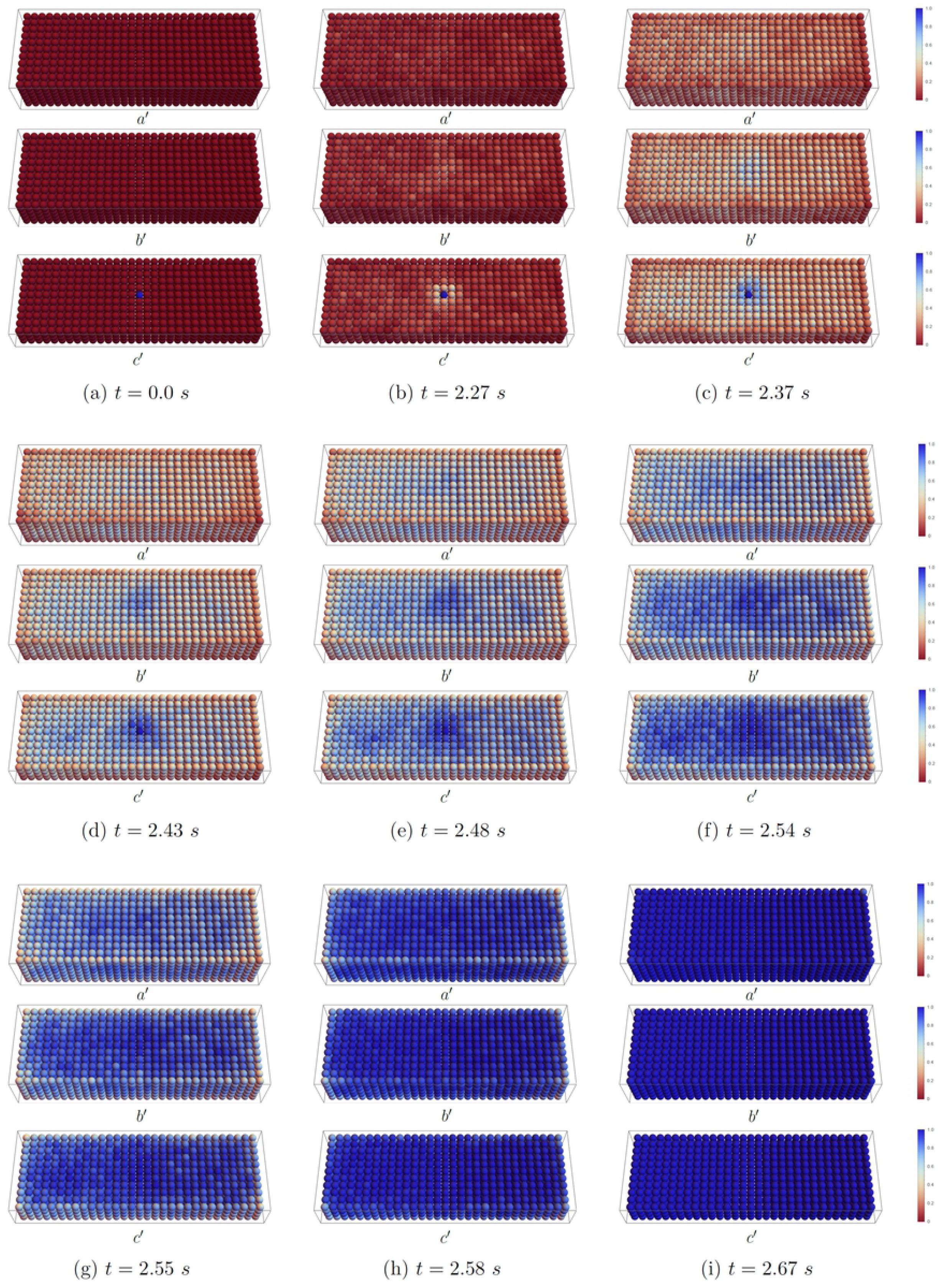
Probability distribution of ice propagation in different time steps of a large tissue with 2387 cells and a pre-frozen cell in the center of the tissue with *α* = 10.4 and *B* = 100 °C/min. The whole tissue is indicated in *a′*, one layer has been removed in *b′*, and two layers have been removed in *c′*.

Our results confirm that the ice propagates most readily in the tissue centre, where the average number of cells connected is higher. In particular, the most rapid propagation in the slab happens with the centre-seeded cell position, with the face-seeded cell position coming in second, and the corner-seeded cell position being the slowest. Moreover, the ice propagation in the face and corner-seeded slabs typically progresses towards the centre of the tissue and then outward as opposed to along an edge or face. This phenomenon reflects the relative connectivity of cells at the centre. In more realistic tissues, the intercellular connection numbers may vary spatially, and our current model is uniquely capable to address this. Note, also, that as the tissue size increases, the average connectivity between cells increases, and depending on *α* the rate of complete ice formation in a well-connected large tissue could be higher than that of a less-well-connected small tissue.

Precision cut liver slices (PCLS) have significant clinical and experimental relevance, but currently do not have an acceptable cryopreservation protocol [58]. PCLS are typically 2-5 mm diameter cylinders of 250 *μ*m thickness [59]. Therefore the current slab model captures between 5% and 1% of the total volume of a 2 to 5 mm diameter PCLS, respectively. Both PhysiCell and our Gillespie Monte Carlo method scale linearly with cell number and both take advantage of OpenMP parallelization. In fact, PhysiCell has been implemented in very high performance computing centres, suggesting that scale up to full size PCLS is well within computational reach, even with the requirement of multiple numerical experiments to inform the Monte Carlo Model [60].

More anatomically realistic implementations of liver tissue using cell-agents can be implemented. For an example, recently Wang *et al*. [55] implemented realistic hepatic lobules using PhysiCell and even account for sources and sinks of transport via the vascular system. Their purpose was tumour modelling, but adaptation to using our model is relatively straightforward. In fact, there is much interest in modeling ice propagation in tissues and organs for the purposes of tumour ablation, known as cryosurgery [61]. Ice propagation in complex tissues is thought to occur first in the vasculature, followed by nucleation in capillaries where it then spreads throughout the tissue or organ [62]. Therefore, the model described by Wang *et al*. is ideally situated to be adapted using our current methods to ice propagation modelling, both from the preservation side, and from the tumour ablation side.

In large tissues, different cooling rates are experienced in different parts of the tissue. In the current manuscript, it was assumed that there were no thermal gradients in the tissues, a reasonable assumption for tissues of diameter of less than 150 *μ*m at the moderate cooling rates studied here. However, the sensitivity of the probability of intracellular ice formation on the temperature of the cell, shown in Fig 8, suggests that in larger tissues such as PCLS or organs accounting for the heat transfer in the system will become important. Moreover, in the current manuscript, the extracellular medium for each cell was assumed to be in a thermodynamic equilibrium that is solely a function of temperature. Anderson, Benson and Kearsley show that there can be a spatial dependence of concentration in the extracellular milieux during cooling [63] on the individual cell level, and Warner *et al*. suggest that, even at superzero temperatures, there is a complicated interaction between solute, solvent, and cellular transport as the size of the tissue increases [64]. Therefore, future work in larger tissues should pay close attention to the possible distribution of both heat and concentration, their effects on the intracellular state of each agent, and in turn, the likelihood that any cell may experience intracellular ice formation. Wang *et al*. used the finite volume based reaction diffusion equation solver associated with PhysiCell to describe the transport of sugars and oxygen for cell support, but this mechanism can be repurposed to capture heat and solute transport relevant to cryobiological applications.

## Conclusions

This study applied a lattice-free agent-based model in combination with a stochastic model for ice formation and propagation in an arbitrary tissue. This method was implemented with the advanced and robust open source package PhysiCell. In agent-based models, cells, as agents, can move and interact with each other freely, and in our implementation, the movement was influenced by the expansion of intracellular water as ice. The probability of ice formation in each cell at a given time is obtained based on two non-dimensional parameters *τ* and *α*. The non-dimensional parameter *τ* in this model, makes it possible to calculate the probability independent of cell type. However, to obtain the real time instead of *τ* which is dependent on the cell type, the information about the mechanism of ice formation inside the cell is needed. Here, we proposed a formulation to compute *τ* in the absence of CPA. We showed that *τ* is a function of cell type and cooling rate. We conclude that there is a threshold value of *α* = 50 for rat hepatocytes, and that increasing value of *α* beyond this threshold does not affect the probability of IIF. However, the actual value of α depends on the intercellular connections in different tissue types. The higher connection between cells facilitate the ice formation due to ice propagative *J*^p^ and reduce the ice formation due to intracellular ice nucleation *J*^i^. This study has also shown that ice is most likely to propagate into the centre of the tissue where there are a higher number of neighbour cells and connections. The proposed model approximation also shows potential for other problems which will be examined in the future, such as ice expansion and propagation in liver tissue with different cell types, mass and heat transport and presence of cryoprotective agents.

## Acknowledgments

Funding for this study was provided by the Saskatchewan Health Research Foundation, and the Canadian Institute for Health Research.

